# Joint Learning of Drug-Drug Combination and Drug-Drug Interaction via Coupled Tensor-Tensor Factorization with Side Information

**DOI:** 10.64898/2026.02.13.705792

**Authors:** Xiaoge Zhang, Zhengyu Fang, Kaiyu Tang, Huiyuan Chen, Jing Li

**Affiliations:** Case Western Reserve University, Cleveland, Ohio, USA

**Keywords:** Machine Learning, Tensor Decomposition, Drug-Drug Interaction Prediction, Drug Combination Therapy

## Abstract

Targeted drug therapies offer a promising approach for treating complex diseases, with combinational drug therapies often employed to enhance therapeutic efficacy. However, unintended drug-drug interactions may undermine treatment outcomes or cause adverse side effects. In this work, we propose a novel joint learning framework for the simultaneous prediction of effective drug combinations and drug-drug interactions, based on coupled tensor-tensor factorization. Specifically, we model drug combination therapies and DDI by representing drug-drug-disease associations and drug-drug interaction profiles as coupled three-way tensors. To address the challenges of data incompleteness and sparsity, the proposed model integrates auxiliary drug similarity information, such as chemical structure similarities, drug-specific side effects, drug target profiles, and drug inhibition data on cancer cell lines, within a multi-view learning framework. For optimization, we adopt a modified Alternating Direction Method of Multipliers (ADMM) algorithm that ensures convergence while enforcing non-negativity constraints. In addition to standard tensor completion tasks, we further evaluate the proposed method under a more realistic *new-drug prediction* setting, where all interactions involving a previously unseen drug are withheld. This scenario closely aligns with real-world applications, in which reliable predictions for emerging or under-studied compounds are essential. We evaluate the proposed method on a comprehensive dataset compiled from multiple sources, including DrugBank, CDCDB, SIDER, and PubChem. Our experiments show that SI-ADMM maintains robust performance and achieves the best results comparing to other tensor factorization approaches, with or without auxiliary information, particularly in the new-drug prediction setting. The implementation of our method is publicly available at: https://github.com/Xiaoge-Zhang/SI-ADMM.

## 1 Introduction

In the treatment of complex diseases such as cancer and neurological disorders, combinational drug therapies have shown promise in improving treatment efficacy by targeting multiple biological pathways simultaneously [37]. However, as the number of available drugs continues to grow, the space of possible drug combinations expands rapidly, increasing the risk of undetected drug-drug interactions (DDIs) [46]. Such interactions can reduce therapeutic effectiveness, induce adverse side effects [17], and in severe cases lead to increased hospitalization [10], posing significant challenges to the screening and optimization of combinational drug therapies. Unexpected DDIs can therefore substantially undermine the benefits of combination treatments, either by diminishing efficacy or by introducing harmful effects. Consequently, there is a critical need for computational methods that can accurately predict both beneficial drug-drug synergies and adverse interactions to support safer and more effective therapeutic strategies. Strictly speaking, drug combination therapies represent one category of drug-drug interactions that rely on synergistic or additive effects to increase efficacy, reduce toxicity, and/or prevent drug resistance. Throughout this paper, we use the terms *drug combination* and *drug-drug synergy* interchangeably to denote positive interactions, while the term *drug-drug interaction* is primarily used to refer to adverse interactions that may cause unexpected side effects.

In recent years, a variety of machine learning and computational approaches have been proposed to predict DDIs and drug-drug synergies [52, 11, 12], with several methods leveraging tensor decomposition or tensor completion techniques [54]. Tensor-based models are particularly attractive because they can naturally integrate multi-relational information involving drugs, targets, and diseases within a unified representation. Despite these advantages, most existing studies treat DDI prediction and drug-drug synergy prediction as separate problems [38, 43], and relatively few efforts have attempted to jointly model these two closely related interaction types within a single computational framework.

The motivation for our study is rooted in the observation that drug-drug synergies and adverse DDIs are essentially two sides of the same coin: while one emphasizes positive therapeutic effects, the other focuses on identifying harmful interactions that may lead to side effects. Although the outcomes differ, these interaction types may share underlying biological and pharmacological mechanisms. Jointly modeling both forms of interactions has the potential to improve predictive accuracy and provide richer insight into interaction effects. However, this problem cannot be reduced to a simple binary classification task that assigns positive or negative labels to drug pairs, as interaction outcomes are highly context-dependent, influenced by factors such as disease specificity and interaction mechanisms.

To address these challenges, we propose a novel joint learning framework based on coupled tensor-tensor decomposition that simultaneously models drug-drug-disease associations and adverse drug-drug interaction profiles. To mitigate the severe sparsity inherent in these data sources, we incorporate multiple forms of auxiliary drug side information—including chemical structures, individual drug side effects, and drug target profiles—within a multi-view learning architecture. Building on this formulation, we develop an efficient optimization algorithm based on the Alternating Direction Method of Multipliers (ADMM) [4, 35, 20, 41, 5], enhanced with structured side information, which we refer to as **SI-ADMM**. We evaluate the proposed method on a comprehensive dataset compiled from Drug-Bank [21], PubChem [19], DrugCombDB [30], SIDER [25], and BindingDB [13]. Experimental results demonstrate that SI-ADMM consistently outperforms existing tensor decomposition methods in predicting both drug-drug synergies and adverse interactions, while ablation studies further confirm its robustness across a wide range of parameter settings.

### The main contributions of this paper are summarized as follows

1. We propose a unified joint learning framework that integrates drug synergy prediction and adverse interaction prediction via coupled tensor-tensor decomposition.
2. We develop SI-ADMM, an enhanced ADMM-based optimization algorithm that incorporates auxiliary information to mitigate data sparsity.
3. We demonstrate the effectiveness of SI-ADMM on large-scale datasets compiled from multiple biomedical databases, achieving competitive performance over state-of-the-art tensor decomposition methods.
4. We conduct comprehensive ablation studies showing the robustness of SI-ADMM, further validating its practical utility.

## 2 Related Work

Tensor decomposition has become an effective and widely used framework for modeling and predicting drug-drug interactions (DDIs) and drug-drug synergies, particularly due to its ability to capture higher-order relationships among multiple biological entities. Classical tensor decomposition techniques, including Tucker decomposition and CAN-DECOMP/PARAFAC (CP) decomposition [22], have been extensively applied to biomedical data analysis. In the context of DDI prediction, these methods are typically used to recover missing entries in partially observed tensors by learning low-rank latent factors that encode interaction patterns among drugs, diseases, or experimental conditions.

Beyond purely latent factor models, a substantial body of work has emphasized the importance of incorporating auxiliary or side information to improve predictive performance and biological interpretability. Side information may include chemical structure similarities, molecular fingerprints, gene expression profiles, protein-protein interaction networks, pathway annotations, or known drug-target associations. Prior studies have consistently shown that integrating such heterogeneous information sources helps mitigate data sparsity and enhances generalization, particularly in cold-start or new-drug scenarios [9, 6]. These approaches align with the guilt-by-association principle [7, 24], where drugs sharing similar biological or chemical characteristics are more likely to exhibit similar interaction behaviors.

Several tensor-based models have explicitly incorporated side information through regularization or constrained factorization. For example, Zhang et al. [52] proposed a constrained tensor factorization (CTF-DDI) model that embeds biological constraints and similarity information into the tensor decomposition process, enabling improved robustness under extreme sparsity. Ding et al. [11] introduced a deep tensor factorization (DTF) framework for anticancer drug synergy prediction, leveraging deep neural architectures to fuse multiple data modalities. While these methods demonstrate strong empirical performance, they typically focus on a single tensor and do not explicitly model interactions across multiple related relational tensors.

In parallel, coupled and joint factorization methods have been explored in broader machine learning and signal processing communities. Joint tensor decomposition and coupled tensor-matrix factorization have shown success in applications such as multi-view learning, recommender systems, and image processing [43]. In the biomedical domain, coupled tensor-matrix decomposition has been applied to tasks such as drug-target interaction prediction and multirelational biological data integration [15]. These methods exploit shared latent factors across different data sources to capture consistent underlying structures while preserving modality-specific information.

Despite these advances, the application of joint tensor decomposition to drug discovery remains relatively limited. In particular, existing DDI and drug synergy models often treat different relational datasets independently, even when they involve the same set of drugs. In this work, we address this gap by jointly decomposing two related tensors: one encoding drug-drug-disease relationships and the other representing drug-drug interaction outcomes. By sharing latent drug factors across both tensors, our approach enables information transfer between complementary tasks, facilitating simultaneous prediction of drug combination efficacy and potential adverse interactions. Furthermore, we incorporate multiple auxiliary similarity matrices into the optimization framework, allowing the model to leverage diverse biological prior knowledge while maintaining a unified and interpretable latent representation.

## 3 Problem Formulation

### 3.1 Prediction as Tensor Recovery

To simultaneously predict drug-drug combinations and adverse drug-drug interaction within their context, we assume that drug-drug-disease relationships are encoded in a three-way *drug* ×*drug* × *disease* tensor *𝒳* of size *n* ×*n* ×*m*, where *n* and *m* represent the number of drugs and the number of diseases, respectively. An entry *𝒳*_*ijk*_ = 1 indicates that drug *i* and drug *j* have a synergistic effect on disease *k*, while entries are set to 0 otherwise.

We assume that drug-drug interactions are encoded in a three-way *drug*× *drug*× *DDI* tensor 𝒴 of size *n* ×*n×t*, where *n* and *t* denote the number of drugs and the number of DDI types, respectively. An entry *𝒴*_*ijk*_ = 1 indicates that drug *i* and drug *j* are reported to DDI *k* when used together, with entries set to 0 otherwise. We only have incomplete observations for both tensors *𝒳* and 𝒴, which are denoted as 𝒯 and 𝒪, respectively. The goal of joint drug-drug combination and drug-drug interaction prediction is to recover *𝒳* and 𝒴 from 𝒯 and 𝒪. We formulate this problem as a coupled tensor-tensor completion problem because the two tensors share the same set of drugs. Throughout the paper, we use calligraphic letters (*𝒳, 𝒴*) to denote tensors, and bold uppercase letters to denote matrices.

### 3.2 Side Information for Data Sparsity

In real-world applications, tensors 𝒯 and 𝒪 are often sparse, with many unknown entries. Recovering these unobserved entries based solely on the tensors is challenging. To address this, we incorporate additional auxiliary information and formulate the problem under the multi-view learning framework. In this work, we primarily focus on side information related to drugs, because the same set of drugs is present in both tensors.

We further assume that all the side information can be organized into similarity/kernel matrices, which are then integrated into the learning process and jointly decomposed with the two tensors. We denote the similarity matrices constructed from *n*_*a*_ views of drugs as 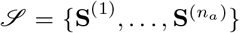. In practice, these similarity matrices are constructed based on different properties of drugs such as their chemical structures and side effect information when drugs are used alone. We will discuss more side information in Section 5. The overall architecture of our proposed approach is illustrated in Fig 1.

**Figure 1.**
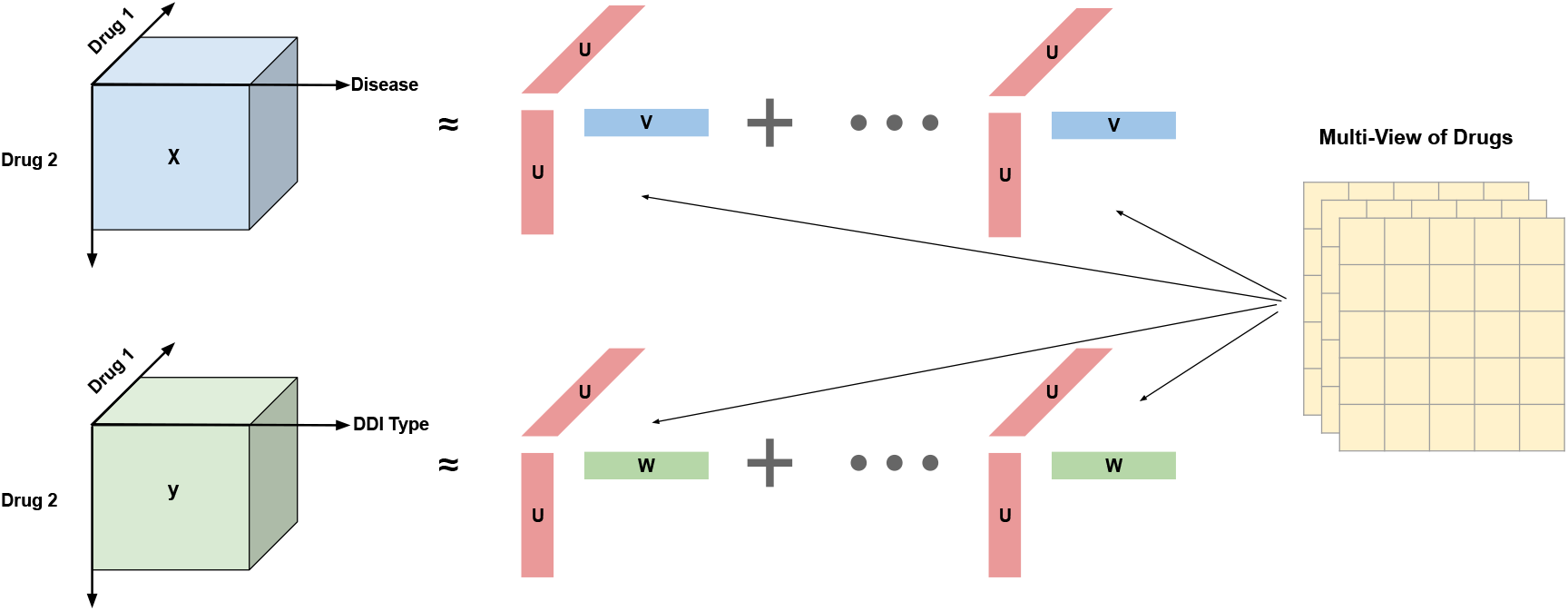
The architecture of SI-ADMM showing the joint tensor-tensor decomposition is aided by auxiliary side information from different views of drugs.

## 4 Methods

### 4.1 Tensor Decomposition

To recover the drug-drug-disease tensor *𝒳* and the DDI tensor 𝒴, we adopt the INDSCAL decomposition method, which is a special case of CP decomposition [22]. Given a three-dimensional tensor ℳ, the goal of CP decomposition is to represent ℳ as a summation of a set of rank one tensors:

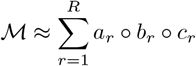

where *R* is a hyperparameter, representing the rank after the decomposition. Vectors *a*_*r*_, *b*_*r*_, *c*_*r*_ are latent factors, and °denote the vector outer product.

In our problem setup, both tensor *𝒳* and tensor 𝒴 have drugs as their first and second mode, which imposes an additional constraint where the latent drug factor in the decomposition must be identical for both tensors, essentially creating a symmetric structure across the drug-drug slices of the tensors. INDSCAL decomposition is precisely for such a structure. More specifically, we want to decompose the two tensors *𝒳* and 𝒴 simultaneously as follows:

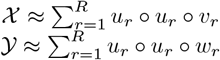

where *u*_*r*_, *v*_*r*_, and *w*_*r*_ represent the drug, disease, and DDI latent factors for the *r*^*th*^ rank component, respectively.

### 4.2 Incorporating Side Information

To overcome the difficulties of decomposing highly sparse tensors, we propose a novel addition to the loss function to incorporate side information of drugs, as mentioned in the problem formulation section. One challenge to directly incorporating similarity matrices from side information is that many of them have significant missing values. To address this challenge, we further decompose each of the similarity matrices into its own latent space and require all lower rank matrices in the latent spaces share some commonality. Finally, the tensor decomposition loss and the side information decomposition loss are combined into one loss function, which is defined as follows:

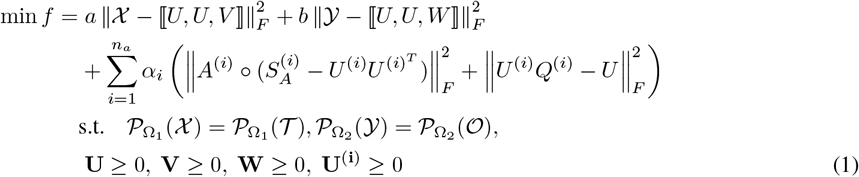

where [**U, U, V**] = Σ _*r*_ *u*_*r*_ ∘ *u*_*r*_ ∘ *v*_*r*_ is a shorthand for the sum of rank-one tensors; **U** is the latent factor matrix of the drug mode for both tensor 𝒳 and tensor 𝒴; **U**^(**i**)^ is the latent drug factor matrix from the *i*-th view of side information; **V** is the latent disease factor matrix for tensor *𝒳*, and **W** is the latent factor matrix for drug-drug interaction tensor 𝒴. *a* and *b* are weight factors which control the weight of the reconstruction losses of *𝒳* and 𝒴 in the case of two tensors having disproportionate sizes. Ω_*i*_ represents the set of observed entries, and 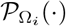 retains the entries in Ω_*i*_ while zeroing out the others [18, 47]. The non-negativity constraints on the latent factors **U, V, W**, and **U**^(**i**)^ help to yield more interpretable and intuitive decomposition results [35, 49].

The last term simultaneously decomposes all drug similarity matrices 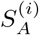into latent factor matrices **U**^(**i**)^, and further requires that **U**^(**i**)^ should be close to the drug latent factor matrix **U** from the tensor decomposition, up to a scaling factor **Q**^(**i**)^. *A*^(*i*)^ is a binary masking matrix for each side of information to mitigate the impact of missing values in the side information. The entry has a value of one for the existing value in the side information matrix and zero otherwise. In this work, we use ∘ to represent Hadamard Product. This last term can also be viewed as an application of guilt-by-association principle [7, 24]. The idea is to use matrix factorization [7, 8, 32] to learn multiple incomplete kernel matrices from multiple data sources representing different views of drugs. Because all the views represent the same entities, they should share some common latent structures. By factorizing all views of drugs into similar latent factors and requiring them to be consistent with the latent factor from the tensors, we can effectively link all the side information together, and link all the side information with the two main tensors representing drug combinations and adverse drug-drug interactions. In addition, by factorizing multiple views simultaneously, this term also mitigates the practical problem of missing data contained in each data source, as suggested by existing work [31, 9]. The purpose of the scaling matrix **Q**^(**i**)^ is to make different views comparable to each other after the decomposition, which is a diagonal matrix that sums over the columns of 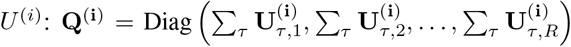. The weight parameter *α*_*i*_ represents the relative importance of the *i*-th view of drugs, *i*.*e*., how influential **S**^(**i**)^ is in shaping the decomposition, and can be determined based on prior knowledge about the side information or through empirical studies.

### 4.3 Learning Algorithm

Many algorithms have been proposed to solve the non-negative tensor decomposition problem [49, 22, 35, 1, 2]. The majority of the optimization algorithms use multiplicative update rules [49] or the gradient descent method [1]. However, the non-negativity constraints may bring a significant computational burden with slow convergence and complicate the development of parallel algorithms to compute large datasets [28]. Also, in such cases, the INDSCAL model is not well understood when the factorization is symmetric [2, 22]. In this work, we adapt the ADMM algorithm to optimize the latent factor matrices based on the partially augmented Lagrangian method. Notice that the objective function in Eq.(1) is non-convex w.r.t. **U, V, W**, and **U**^(**i**)^ all together and it involves fourth-order terms w.r.t. **U** and **U**^(**i**)^, which is hard to optimize directly. To mitigate such problems, SI-ADMM consists of the following steps. First, it converts the objective function to a function of a lower order by using the variable splitting technique. Second, to solve the optimization problem with constraints, we rely on the augmented Lagrangian method, and the new objective function is derived. The third step consists of the core of ADMM, which updates one variable while fixing the others. The details of these steps are discussed in the sequel.

#### Algorithm 1 Pseudo-code of SI-ADMM

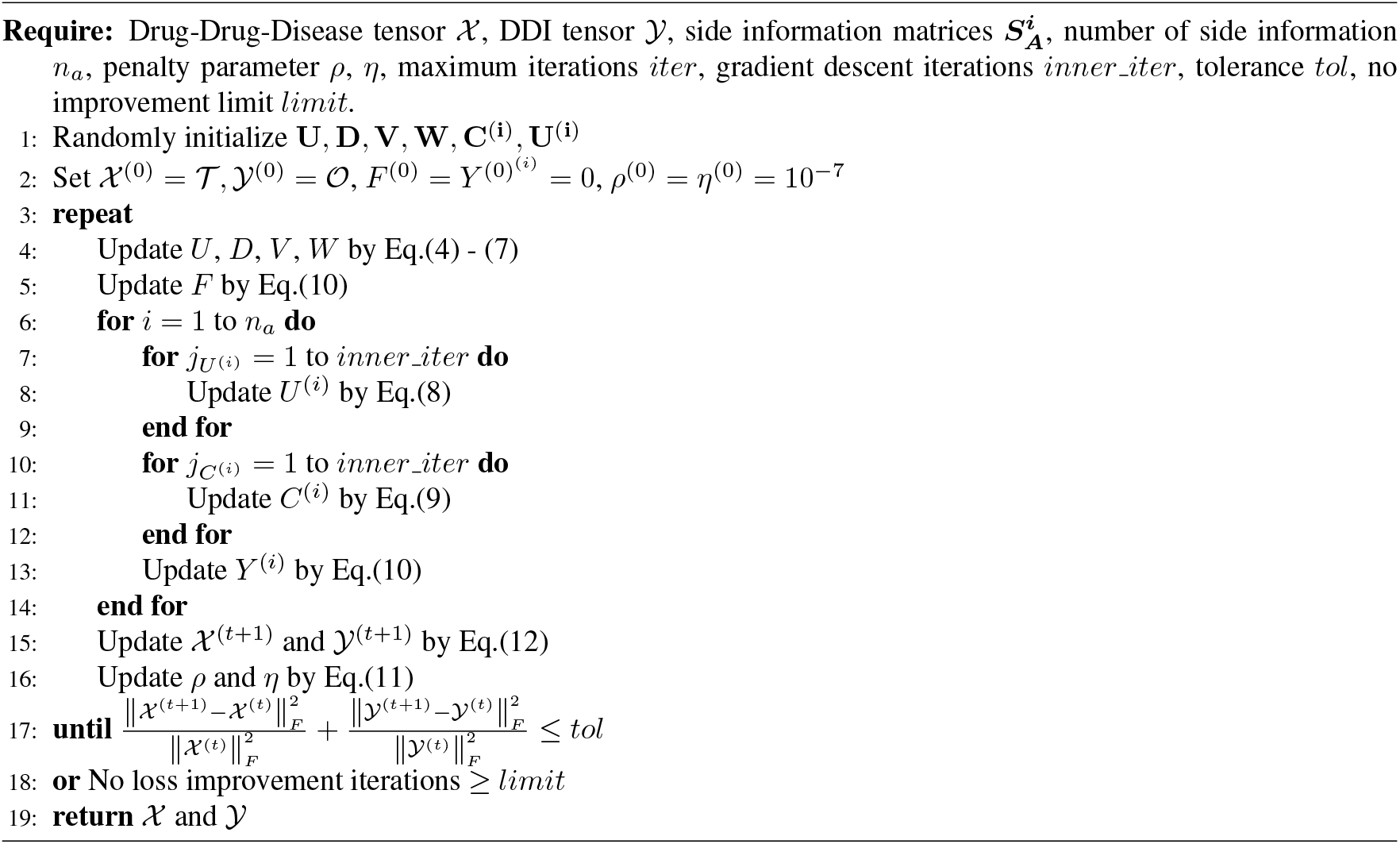

#### 4.3.1 Lower Order Function Using Variable Splitting

To use the variable splitting technique, we define two sets of new variables **D** and **C**^(*i*)^ to replace some occurrences of **U** and **U**^(*i*)^ in Eq.(1), and simultaneously add the constraints **D** = **U** and **C**^(*i*)^ = **U**^(*i*)^ to the system, which enables the reduction of the order of the objective function from 4 to 2 and makes the variables more separable for the ADMM algorithm. The new objective function becomes:

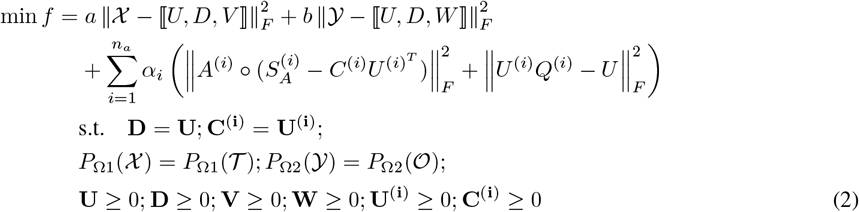

#### 4.3.2 Partially Augmented Lagrangian Function

To utilize the ADMM algorithm to solve the optimization problem, we form the following partially augmented Lagrangian function ℒ :

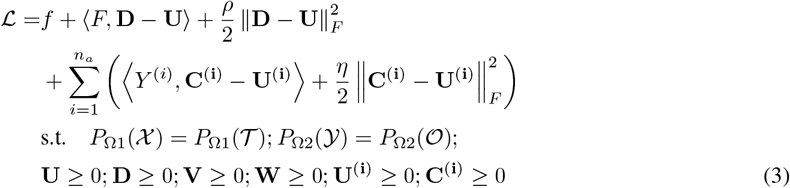

where *f* is the objective function in Eq.(2). The notation ⟨*A, B*⟩denotes the inner product of two matrices. The additional linear order terms are added to handle the constraints **U** = **D** and **C**^(*i*)^ = **U**^(*i*)^, where *F* and *Y* ^(*i*)^ in these terms are Lagrange multiplier. The additional second-order terms are for regularization. Penalty parameters *ρ* and *η* are applied to the constraint of *U* = *D* and side information constraints of *C*^(*i*)^ = *U* ^(*i*)^, respectively.

#### 4.3.3 The SI-ADMM Algorithm

Following the original ADMM framework, we adopt an alternating optimization strategy, where each latent matrix is updated sequentially while keeping the others fixed until convergence is achieved. The update process for each matrix consists of three key steps:

1. **Term Identification:** Extract all terms from the partial augmented Lagrangian that depend on the matrix to be updated.

2. **Derivative Computation:** Calculate the partial derivative with respect to the target matrix, and derive the closed-form solution by setting the derivative to zero.

3. **Matrix Update:** Apply the derived update rule and use the updated matrix in the subsequent optimization steps.

We present full derivations for the updates of *U*, the Lagrangian multipliers, penalty terms, and associated tensor components in the following section. Derivations for the remaining latent matrices are provided in the supplementary appendix.

1. **Update of** *U* The terms in the Lagrangian ℒ involving *U* are expressed as:

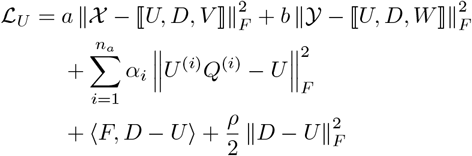

Since the direct computation of the tensor reconstruction norm is non-trivial, we leverage the matricization and Khatri-Rao product approximation introduced in [22], where the tensor decomposition can be written as:

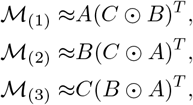

Here, ℳ is a 3-D tensor and *A, B, C* are the latent factor matrices. ℳ_(1)_, ℳ_(2)_, ℳ_(3)_ are the mode-1, mode-2, and mode-3 matricization of the tensor [22]. ⊙ is Khatri-Rao Product. Using this approximation rule, we can rewrite the terms involving *U* into:

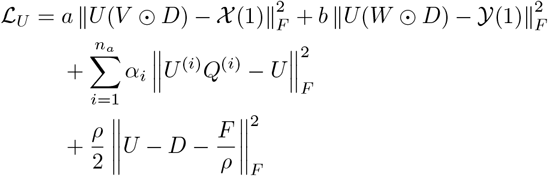

Defining Π = *V* ^(*t*)^ ⊙*D*^(*t*)^ and Θ = *W* ^(*t*)^ ⊙*D*^(*t*)^, and following the matrix derivative identities provided in [36], the partial derivative with respect to *U* becomes:

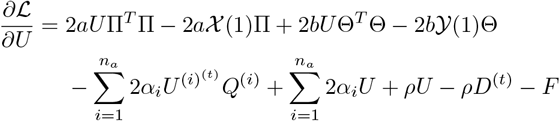

Here *t* is the *t*th iteration of optimization. By equating the derivative to zero, the update rule for *U* is derived as:

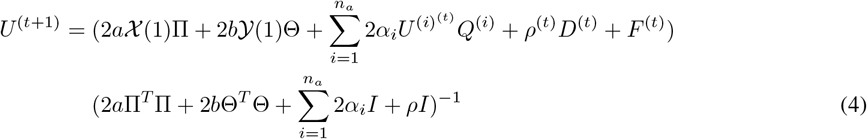
2. **Update of Lagrangian Multipliers** Unlike standard ADMM, which updates the Lagrangian multipliers for all imposed constraints, our framework updates the multipliers specifically for the variable splitting constraints, namely *U* = *D* and *U* ^(*i*)^ = *C*^(*i*)^. The updates are performed as follows:

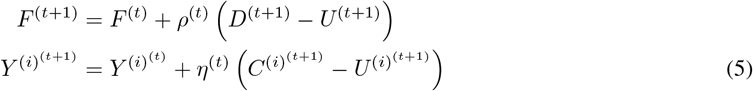
3. **Update of Penalty Parameters** To enhance convergence speed, we follow the adaptive penalty update strategy proposed in [29], updating the penalty parameters as:

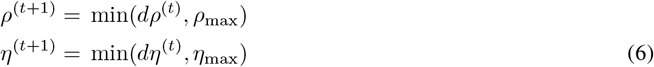

where *d* is a scalar larger than 1, and *ρ*_max_ and *η*_max_ are upper bounds to prevent the penalties from becoming excessively large.
4. **Update of Predicted Tensors** At each iteration, the unobserved entries of the tensors are estimated by combining the observed data with the reconstructed values from the current latent factors, according to the following update rules:

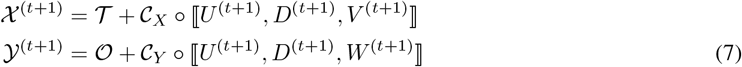

Here, *𝒳*^(*t*+1)^ and 𝒴^(*t*+1)^ represent the predicted tensors at iteration *t* + 1, while 𝒯 and 𝒪 denote the observed parts of the tensors, as defined in Section 3.1. The complement masks 𝒞_*X*_ and 𝒞_*Y*_ have the same dimensions as 𝒯 and 𝒪, respectively, containing zeros at observed entries and ones at unobserved entries. ° means element-wise product.
5. **Summary of the SI-ADMM Algorithm** Combining all the update procedures described above, as well as the initialization and termination steps, the complete SI-ADMM algorithm is summarized in Algorithm 1. For the termination step, we check the weighted reconstruction loss of both tensors*𝒳* and𝒴 against a tolerance threshold at the end of each iteration (line 12 in Algorithm 1), and the algorithm stops when the difference between two consecutive runs is smaller than the threshold or the number of iterations with no improvement in reconstruction loss reaches the limit.

### 4.4 Complexity Analysis For SI-ADMM

The main computational cost arises from updating the matrices *U, D, V, W, U* ^(*i*)^, and *C*^(*i*)^. The most expensive terms in these updates are as follows. First, the term *𝒳*_(1)_*V* ⊙*D* + 𝒴 _(1)_*W* ⊙ *D*, where⊙ denotes the Khatri-Rao product, requires 𝒪 (*N* ^2^*M*_disease_*R* + *N* ^2^*M*_DDI_*R*) operations, with *N* the number of drugs, *M*_disease_ the number of diseases, and *M*_DDI_ the number of DDI types. Second, the term (*V* ⊙*D*)^T^(*V* ⊙*D*) + (*W*⊙ *D*)^T^(*W* ⊙*D*) can be equivalently written as *V* ^T^*V D*^T^*D* + *W* ^T^*WD*^T^*D*, with complexity 𝒪 ((*N* + *M*_disease_)*R*^2^ +(*N* + *M*_DDI_)*R*^2^). Third, computing the matrix inverse required when separating *U, D, V*, and *W* involves a Cholesky decomposition, which has complexity 𝒪 (*R*^3^). Finally, solving the update rule as a whole requires 𝒪(*NR*^2^). Therefore, the overall complexity of SI-ADMM is

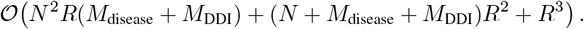

Although the rank *R* appears in higher-order terms, in practice it is much smaller than the number of drugs *N* or (*M*_disease_ + *M*_DDI_).

## 5 EXPERIMENTS

### 5.1 Dataset Preparation

#### 5.1.1 Data Preparation for *𝒳*and 𝒴 tensor

To construct the drug-drug combination tensor and DDI tensor, we curated a task-specific dataset based on two primary sources: CDCDB^1^ [39] and DrugBank^2^ [21]. CDCDB is a comprehensive repository that integrates drug combination information from multiple resources, including ClinicalTrials.gov^3^ [51], international patents, and the FDA Orange Book^4^ [45]. In this work, we focused on drug combinations involving exactly two distinct drugs, extracted from both patents and ClinicalTrials.gov. We did not include combinations from the FDA Orange Book because they did not specify medical conditions associated with drug combinations.

In total, we collect 12033 unique drug-drug-disease triplets that involve 2652 unique drugs and 10156 unique medical conditions. The fine-grained classification of medical conditions results in an extreme sparse tensor and makes it practically infeasible to predict any meaningful drug combinations for any diseases. To address this, we map the disease conditions to the second level disease classifications based on the International Classification of Diseases (ICD) hierarchy^5^ [50]. This reduces the number of diseases down to 238.

We construct the DDI tensor from the DDI dataset of the DrugBank database^6^, which contains manually curated drug-drug interactions from drug labels as well as primary literature. DrugBank provides 86 types of DDI among drug pairs that cover 1706 unique drugs. Using both datasets and cross-referencing various identifiers, we obtain our own dataset consisting of 1070 unique drugs, 238 different disease types, and 81 types of DDI. Therefore the dimensions of 𝒳 and 𝒴 tensors are (1070, 1070, 238) and (1070, 1070, 81), respectively. Both tensors are very sparse. The percentages of entries that equal 1 are 0.0025% and 0.106% for *𝒳* and 𝒴 tensors, respectively.

#### 5.1.2 Formulation of Side Information matrices

We obtained side information similarity matrices from four different sources: the Tainimoto similarity [44] based on SMILE[48] strings, the cosine similarities among the side effect vectors, the global alignment scores of drugs’ target sequences, and cosine similarities among the drugs’ inhibition scores (IC50) on NCI-60 human cancer cell lines [34]. A detailed description of how we formed such side information matrices is provided below.

**Tanimoto scores** are derived from the SMILES representations of all drugs. We employ the RDKit [26] Python library to compute the Tanimoto similarity between all pairs of drugs, resulting in a complete 1070 × 1070 drug-drug similarity matrix without missing.

**Side Effect Information** is obtained from the SIDER database [25]. Only 776 drugs out of 1070 drugs have side effect information. For each drug, we construct an *n*-dimensional binary vector, where *n* represents the total number of side effects recorded in SIDER. An entry in the vector is set to 1 if the drug is associated with the corresponding side effect, and 0 otherwise. We then compute the pairwise cosine similarity between these vectors to generate the drug-drug similarity matrix.

**Target Sequence Alignment scores** are derived from the target protein sequences associated with each drug in the DrugBank dataset. Out of 1070 drugs, only 890 drugs have their target sequences. For every pair of drugs with target sequences, we apply the Smith-Waterman alignment algorithm [40] to compute the scores between their target sequences. The self-alignment scores of target sequences are then normalized to produce a drug-drug similarity matrix.

**IC**_**50**_ **scores** are obtained from experimental data provided by the NCI-60 project [34]. Out of 1070 drugs, only 235 drugs have IC50 data. For each such drug, we construct a 60-dimensional vector representing its IC_50_ values (a measure of inhibitory potency) across the 60 cancer cell lines. Pairwise cosine similarity is then computed between these vectors to generate the drug-drug similarity matrix.

### 5.2 Experiment Settings

#### 5.2.1 Random Prediction Task

To ensure a fair evaluation, we adopt the following data splitting procedure. Specifically, to simulate partially observed tensors 𝒪 and 𝒯 from the fully observed tensors*𝒳* and 𝒴, we randomly mask out 10% of the entries with value 1. These masked entries serve as the positive samples in the testing set. An equal number of randomly selected zero-valued entries are included as negative samples in the testing set. Once the testing set is defined, the resulting partially observed tensors are used as input to the learning algorithm.

We evaluate the performance using two main metrics - the Area Under the ROC Curve (AUC) and Area Under the Precision-Recall Curve (AUPR) - but also include other metrics such as F1-score, accuracy, recall, specificity, and precision. For all baseline methods and SI-ADMM, we perform five independent runs with different random seeds and report the average performance along with the standard deviation.

#### 5.2.2 New-Drug Prediction Task

To better reflect real-world usage, we also evaluate SI-ADMM and the baselines in a new–drug prediction setting. In this scenario, the model must infer potential drug-drug combinations or DDIs for a previously unseen drug, where all of its interaction entries in the tested tensor are masked. Thus, the model can rely only on side information matrices and cross-tensor signals (e.g., drug-drug combination information when predicting DDIs, and vice versa).

Formally, given a target drug and a tensor (either the DDI tensor or the drug-drug combination tensor), all entries whose row or column correspond to that drug are set to zero and treated as test entries. Under this setting, the CP baseline–which incorporates no side information–cannot produce meaningful predictions. Its ALS updates are described in [22], and when all entries for a drug are masked, the updates return zero vectors. Therefore, we compare SI-ADMM only with TDRC, TFAI, and CTF for this task.

We select the top 10 most interacted drugs from each tensor (based on the number of known interactions) for evaluation, yielding 20 drugs in total. Each drug is tested individually under the new–drug protocol for SI-ADMM and the baselines.

Performance is assessed using hit rate, which is most appropriate for this ranking-based retrieval problem. Since all true interactions involving the target drug are masked, metrics such as RMSE, MAE, accuracy, and AUROC become uninformative due to extreme class imbalance and the dominance of zeros. Hit rate instead measures whether the model ranks true interacting partners within the top-*K* predictions, aligning with practical drug discovery workflows.

### 5.3 Baseline Approaches

To the best of our knowledge, no existing approach adopts the same formulation as the one proposed in SI-ADMM. Consequently, a direct comparison with methods performing the exact same analysis is not feasible. Instead, we compare our method against baseline tensor decomposition approaches that independently decompose individual 3-D tensors, either with or without incorporating additional side information. These baseline methods are described in the following sections.

- **CP**: CANDECOMP/PARAFAC (CP) [22] is a standard tensor decomposition method (see Section 4.1). We use the implementation provided by the TensorLy Python library [23], which optimizes the latent factor matrices via Alternating Least Squares (ALS). CP does not incorporate side information and serves as a fundamental baseline.
- **Tensor Factorization using Auxiliary Information (TF-AI)**: TF-AI [33] incorporates symmetric side information matrices for each mode of the target tensor to guide the decomposition process. It introduces two regularization strategies, tailored for dense and sparse tensors respectively. We include TF-AI as an early method that integrates side information in the form of similarity matrices.
- **TDRC**: TDRC [16] is a tensor decomposition method originally proposed for miRNA-disease-type prediction. It incorporates side information as relational constraints in the optimization objective. We include TDRC as a baseline that leverages side information differently from our SI-ADMM framework, focusing on relational regularization.
- **CTF-DDI**: CTF-DDI [14] is a recent tensor decomposition method designed for DDI prediction, incorporating side information directly into its optimization framework. CTF-DDI integrates two types of side information—structural and biological similarities—by averaging their values. We consider CTF-DDI as a strong and competitive baseline.

### 5.4 Hyper-parameters Settings

#### 5.4.1 Hyper-parameters Tuning for SI-ADMM

Our model consists of several hyper-parameters, including the rank, initial values of penalties (*ρ* and *η*), and weights for the main tensors (*a* and *b*) and side information (*α*_*i*_). Rank is set to 60 by running and evaluating the algorithm using the following rank numbers: {10, 20, 30, 40, 50, 60}, the penalty is set to 10^™^7 with a 15% increase every iteration and is capped at 10^5^. The penalties are used solely to accelerate the convergence of the algorithm, so adjustment of this doesn’t affect the outcome of the program. For weighting the loss contributions of the tensors and side information matrices, we apply a linear scaling based on the inverse of their total magnitude. Specifically, the weights for each tensor or matrix are:

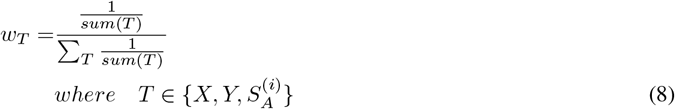

where *sum*(*T*) denotes the sum of all elements, since tensors and side information matrices are all non-negative in *T*. The resulting *w*_*T*_ values serve as the weights (i.e., *a, b*, and *α*_*i*_) in the SI-ADMM optimization framework, balancing the contributions of each tensor and side information matrix. For the dataset used in the SI-ADMM, the weights *a, b*, and *α*_*i*_ are [0.5764 0.04189 0.08469 0.09330 0.02414 0.1795]. For the inner update iterations that updates *U* ^(*i*)^ and *C*^(*i*)^, We set the inner iteration number to 10 after testing the program with five different random seeds across the following inner update iteration counts: 10, 20, 30, 40, 50, 60, 70, 80.

#### 5.4.2 Hyper-parameter Tuning for Baselines

For CTF-DDI [14], we followed the hyperparameter settings reported in their original paper and utilized Tanimoto similarity and target sequence alignments similarity as structural side information, IC50 similarity and side effect similarity as biological side information. For other baseline methods, the primary parameter was the rank number, and we evaluated different rank numbers in {10, 20, 30, 40, 50, 60} and reported their best performance, averaging the metrics over both tensor-recovery tasks.

## 6 RESULTS

### 6.1 Results on Random Prediction Task

Tables 1 and 2 present the prediction results for the drug-drug combination prediction task and the DDI prediction task, respectively, for baseline models and SI-ADMM. The results demonstrate that SI-ADMM consistently achieves the highest performance among all the tested methods in terms of AUC and AUPR (Fig 2), as well as specificity and precision for both the drug-drug combination and DDI prediction tasks. Notably, for the drug-drug-disease association prediction task, SI-ADMM achieves significant improvements in AUPR, AUC, and Precision (all p-values are smaller than 0.05 comparing with the second best method based on the paired t-test). It also achieves the highest score in accuracy and the second highest in F1 score that is nearly indistinguishable from the highest score (from CTF-DDI). For DDI prediction, SI-ADMM is the best approach and achieves the best score in four measures including AUPR and AUC (Fig 2). TDRF is the close second and achieves the best score in the other three measures. These findings highlight the competitive discriminative capability of SI-ADMM in both drug-drug combination and DDI prediction tasks.

**Table 1:**
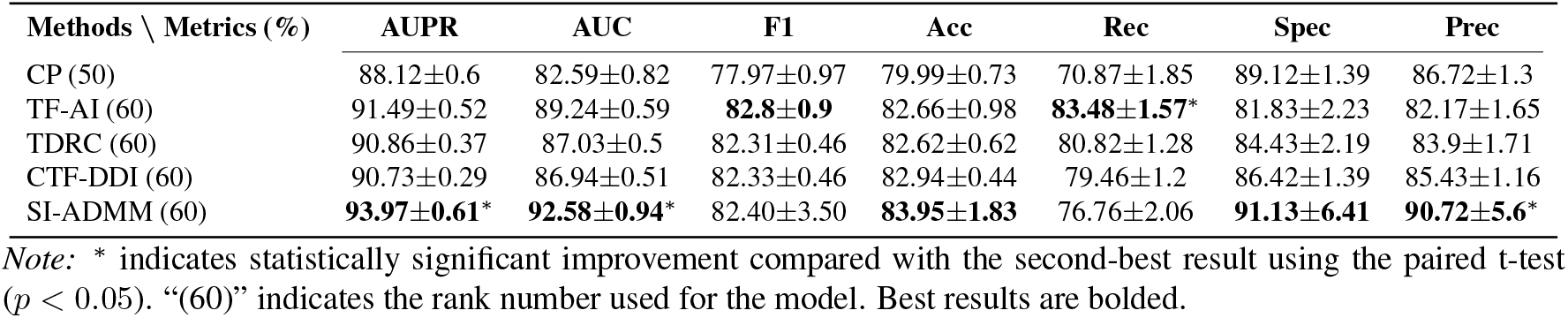
Results of SI-ADMM and baselines on tensor *𝒳* prediction (drug-drug combination)

**Table 2:** Results of SI-ADMM and baselines on tensor 𝒴 prediction (DDI)

| Methods \ Metrics (%) | AUPR | AUC | F1 | Acc | Rec | Spec | Prec |
| --- | --- | --- | --- | --- | --- | --- | --- |
| CP (50) | 98.30±0.08 | 97.50±0.10 | 91.99±0.16 | 92.21±0.17 | 89.41±0.61 | 95.02±0.70 | 94.73±0.67 |
| TF-AI (60) | 97.51±0.03 | 97.66±0.04 | 94.16±0.14 | 94.19±0.13 | **93.63±0.56** | 94.75±0.51 | 94.70±0.47 |
| TDRC (60) | 98.48±0.06 | 97.85±0.09 | **94.34±0.18** | **94.41±0.18** | 93.21±0.29 | 95.61±0.40 | 95.50±0.38 |
| CTF-DDI (60) | 98.48±0.04 | 97.85±0.07 | 94.29±0.10 | 94.40±0.11 | 92.53±0.36 | 96.27±0.52 | 96.13±0.51 |
| SI-ADMM (60) | **98.53±0.06** | **98.21±0.09\*** | 93.12±0.55 | 93.42±0.46 | 89.10±1.56 | **97.74±0.84\*** | **97.55±0.86\*** |

Note: \* indicates statistically significant improvement compared with the second-best result using the paired t-test ( $p < 0.05$ ). “(60)” indicates the rank number used for the model. Best results are bolded.

**Figure 2.**
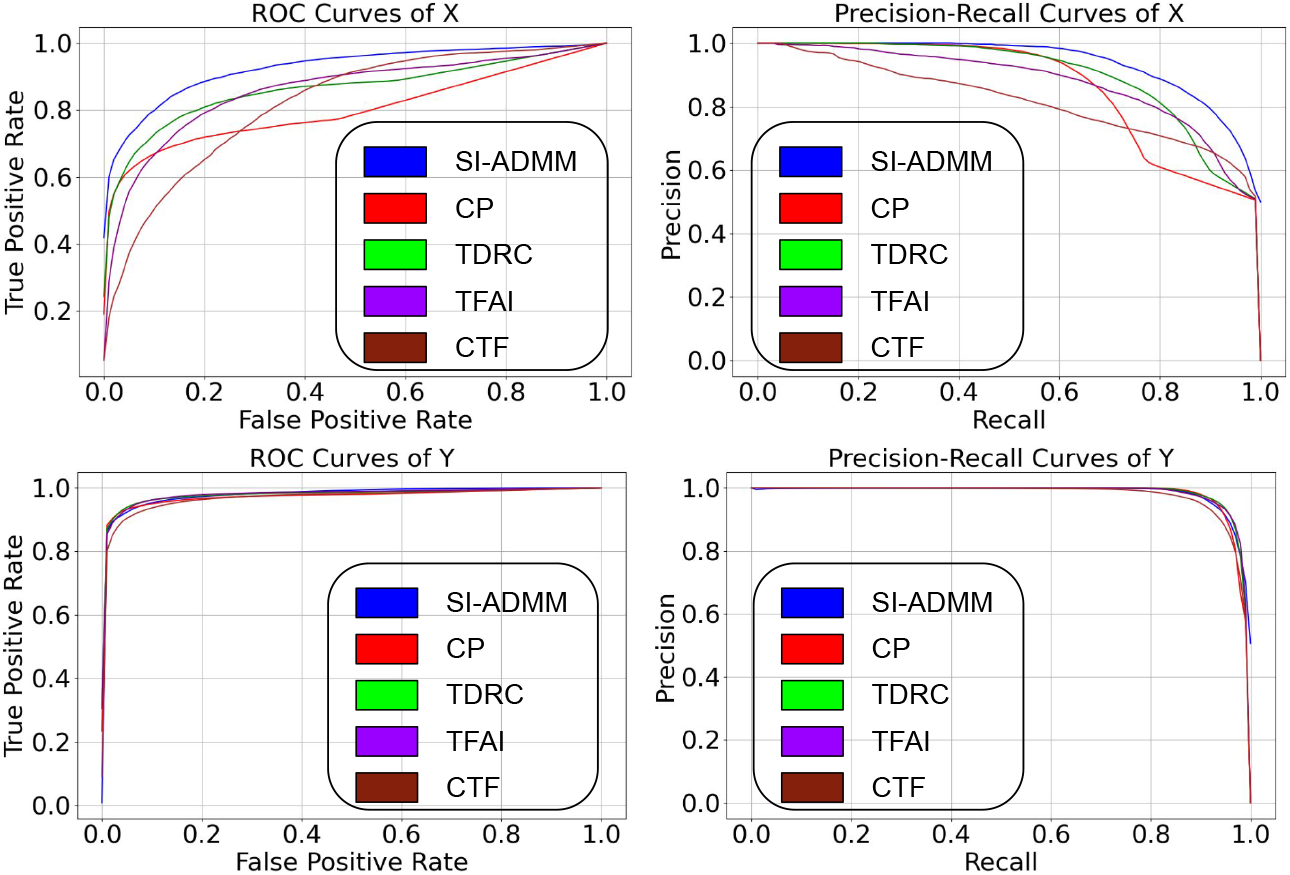
ROC and Precision-Recall curves for different models on tensors *𝒳* (top) and 𝒴 (bottom).

It is also worth noting that all baseline methods perform independent decompositions for each tensor, resulting in separate sets of three latent factor matrices for each task. In contrast, SI-ADMM jointly decomposes both tensors using a shared set of four latent factor matrices, namely *U, D, V*, and *W*. The superior performance of SI-ADMM not only demonstrates that the shared drug latent factors can effectively capture meaningful representations for both tasks but also highlights the model’s strong learning capability despite operating with fewer degrees of freedom in terms of the number of latent factor matrices.

### 6.2 Further Analysis of SI-ADMM

To further analyze the structural properties of the proposed SI-ADMM model, we examine the correspondence between the original tensors *𝒳* and 𝒴 and their predicted counterparts at a global interaction level. For each tensor, interactions are aggregated by summing over the third mode (i.e., the drug-drug-disease or DDI dimension), yielding a symmetric drug-drug matrix in which each entry represents the overall interaction strength between a pair of drugs. We then compute row (equivalently, column) sums of the aggregated matrix and rank drugs according to their interaction intensities. To ensure a consistent basis for comparison, the predicted tensors are aggregated and visualized using the same drug ranking derived from the original tensors. The resulting ranked heatmaps, shown in Figure 3, facilitate a direct comparison of large-scale interaction structures between the original and reconstructed tensors. As observed in the figure, the predicted heatmaps closely match their original counterparts in terms of overall intensity distributions and prominent structural patterns, including dense interaction blocks and banded regions, indicating that SI-ADMM preserves the global organization of drug-drug interactions rather than overfitting individual tensor entries.

**Figure 3.**
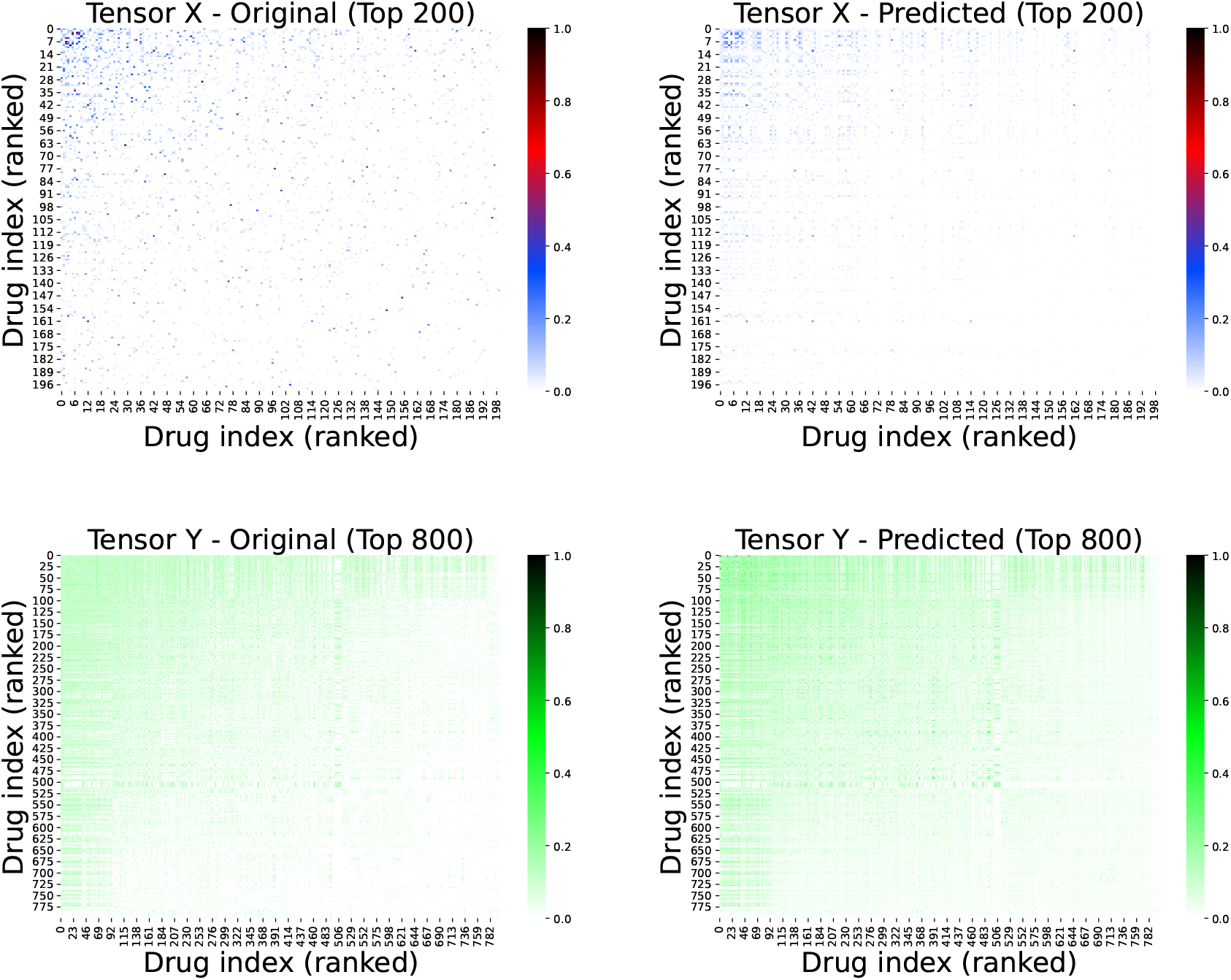
Ranked heatmaps of aggregated tensors. Tensors are summed over the third mode and reordered by row-sum ranking from the original data, which is consistently applied to predictions. The predicted heatmaps recover key block and band structures, demonstrating preservation of global interaction patterns. Owing to sparsity, the top 200 drugs for *𝒳* and top 800 drugs for 𝒴 are shown.

To complement this structural visualization, we quantitatively assess the agreement between the original tensors and their predictions using global similarity metrics. Direct entry-wise comparison is unreliable due to the extreme sparsity of the original tensors; therefore, all quantitative evaluations are conducted on the aggregated two-dimensional interaction matrices described above. This aggregation retains meaningful global interaction structure while reducing sparsity-induced noise, enabling a more stable and informative assessment. Specifically, we compute Pearson correlation [3], Spearman rank correlation [42], and relative Frobenius error between the aggregated original and predicted matrices. These measures respectively capture linear correspondence, rank-level consistency, and overall reconstruction fidelity, and are commonly used to evaluate structural similarity in high-dimensional matrix and tensor analysis.

The quantitative results demonstrate a strong structural correspondence for tensor 𝒴, with high Pearson (0.79) and Spearman (0.59) correlations and a low relative Frobenius error (0.65), suggesting that SI-ADMM effectively recovers dominant drug-drug interaction patterns. For tensor *𝒳*, a moderate Pearson correlation (approximately 0.46) indicates that the model captures coarse interaction intensity structure, while weaker rank-level agreement reflects heterogeneity across disease-specific tensor slices. In contrast, entry-wise correlations computed directly on the full three-dimensional tensors are close to zero, which is expected given the high dimensionality and sparsity of biomedical interaction data and does not contradict the model’s ability to recover meaningful global structure. Taken together, these results provide evidence that SI-ADMM captures latent interaction patterns in a structurally consistent manner at an analytically relevant scale.

### 6.3 Results on New-Drug Prediction Task

Table 5 summarizes the results of New-Drug Prediction for SI-ADMM and the applicable baselines measured using hit rate. SI-ADMM consistently achieves strong hit-rate performance across the most interacted drugs for both tensors, with particularly notable gains on the 𝒴 tensor. These results demonstrate that the proposed joint learning framework can effectively exploit side information and cross-tensor coupling to infer meaningful interaction patterns even in the absence of historical interaction data. Compared to competing baselines, the consistently higher hit rates indicate that SI-ADMM generalizes more reliably under cold-start conditions, which closely reflect real-world deployment scenarios where newly developed or insufficiently studied drugs are common. Consequently, the strong performance advantage observed in this new-drug setting suggests that SI-ADMM has a higher likelihood of delivering improved predictive accuracy in practical applications, where robust interaction prediction based on limited information is essential.

### 6.4 Case Analysis

After running the algorithm, we examined the entries with the top 10 highest predicted values for both tensors. The top 10 results for tensor *𝒳* are shown in Table 3, where the drug pairs are represented using their DrugBank IDs. We observe that although our model does not directly impose symmetric constraints on the drug pairs in the recovered tensors, the prediction results show many drug pairs (8 out of 10) of the top-ranked predictions are symmetric entries. For example, the prediction values of entry (Ibuprofen/DB01050, Acetaminophen/DB00316) and entry (Acetaminophen/DB00316, Ibuprofen/DB01050) ranked top 2 and they were for the same disease. This is also true for tensor 𝒴, in which case the top 10 predictions comprise five unique drug pairs.

**Table 3:** Top-10 predictions from tensor *𝒳*.

| Rank | Drug Pair | Disease |
| --- | --- | --- |
| 1-2 | (Docetaxel/DB01248, Cisplatin/DB00515) | Malignant neoplasms |
| 3-4 | (Cisplatin/DB00515, Gemcitabine/DB00441) | Malignant neoplasms |
| 5 | (Docetaxel/DB01248, Doxorubicin/DB00997) | Malignant neoplasms |
| 6-7 | (Irinotecan/DB00762, Cisplatin/DB00515) | Malignant neoplasms |
| 8 | (Ibuprofen/DB01050, Acetaminophen/DB00316) | Arthropathies |
| 9-10 | (Lenalidomide/DB00480, Bortezomib/DB00188) | Neoplasms |

Note: Drug pairs are unordered; inverse pairs may appear at lower ranks in the full prediction list.

Furthermore, the suggested drug combinations are clinically plausible based on their known functions and descriptions. For example, Ibuprofen/DB01050 and Acetaminophen/DB00316 are two commonly used over the counter medicines for pain and inflammation management. Their combination aligns well with their predicted disease association (arthropathies). All other drugs (Docetaxel/DB01248, Cisplatin/DB00515, Gemcitabine/DB00441, Doxorubicin/DB00997, Irinotecan/DB00762, Lenalidomide/DB00480, and Bortezomib/DB00188) are commonly used in cancer treatment, supporting their predicted links to various neoplasms. In terms of the DDI prediction, the top 10 predictions are composed by five symmetric pairs and is shown in Table 4. 4 out of 5 pairs involves in the same DDI type: *one drug may increase the anticoagulant activities of the other drug*. The drugs in these four pairs (Acenocoumarol/DB01418, Warfarin/DB00682, Phenprocoumon/DB00946, Bivalirudin/DB00006, Rivaroxaban/DB06228) are all blood thinner, which supports the predicted DDI type.

**Table 4:** Top-10 predictions from tensor 𝒴.

| Rank | Drug Pair | DDI Type |
| --- | --- | --- |
| 1-2 | (Acenocoumarol/DB01418, Warfarin/DB00682) | Increasing anticoagulant activities |
| 3-4 | (Acenocoumarol/DB01418, Phenprocoumon/DB00946) | Increasing anticoagulant activities |
| 5-8 | (Avanafil/DB06237, DB00484) | Increasing antihypertensive activities |
| 6-7 | (Warfarin/DB00682, Bivalirudin/DB00006) | Increasing anticoagulant activities |
| 9-10 | (Rivaroxaban/DB06228, Bivalirudin/DB00006) | Increasing anticoagulant activities |

*Note:* Drug pairs are unordered; inverse pairs may appear at lower ranks in the full prediction list.

**Table 5:** Results of SI-ADMM and baselines on New-Drug prediction (top-10 most interacted drugs).

| Method | HR@100 $\mathcal{X}$ | HR@100 $\mathcal{Y}$ |
| --- | --- | --- |
| TF-AI | 10.00±3.79 | 24.40±7.20 |
| TDRC | 17.60±8.80 | 29.00±10.04 |
| CTF-DDI | 22.00±11.73 | 33.60±10.50 |
| SI-ADMM | **29.80±17.18** | **55.20±16.50\*** |

*Note:* \* indicates statistically significant improvement compared with the second-best result using the paired t-test ( $p < 0.05$ ). Best results are bolded.

all the 5 unique drug pairs in the top 10 predictions involve a common drug, Hydroxyzine/DB00557 and all drug pairs are associated with the same DDI type: *one drug may increase the central nervous system (CNS) depressant activities of the other drug*. Hydroxyzine is a prescription medication used to treat anxiety and tension, while its pairing drugs—Buprenorphine/DB00921, Tapentadol/ DB06204, Zolpidem/DB00425, Droperidol/DB00450, and Thalidomide/DB01041 — are known to act as painkillers, neural suppressants, or anti-inflammatory drug. This supports the plausibility of the predicted interactions, as combining these drugs could indeed potentiate CNS depressant effects.

### 6.5 Ablation Studies

To show the robustness of the model, we carried out the following ablation studies: the impact of tensor weights (*a* and *b*) on the tensor reconstruction losses and the impact of the rank parameter on performance of all the approaches.

#### 6.5.1 Impact of Tensor Weights on Reconstruction Losses

To evaluate the effectiveness of the weight selection in Eq. (13), we varied the tensor weights *a* and *b* for *𝒳* and 𝒴 individually while fixing all other weights, and compared the resulting reconstruction losses on the test indices between the original and predicted tensors. As shown in Figure 4, increasing *a* or *b* reduces the reconstruction loss for the corresponding tensor (*𝒳* or 𝒴), but does so at the expense of a significant increase in the loss of the other tensor whose weight remains unchanged. The weights specified in Eq. (13) therefore achieve the best trade-off, minimizing the overall reconstruction loss across both tensors. We also observe that the reconstruction loss of an individual tensor (e.g., *𝒳*) does not necessarily decrease monotonically during optimization, which can be attributed to the fact that the overall objective function consists of multiple coupled terms, and the loss associated with *𝒳* represents only one component of the joint optimization objective.

**Figure 4.**
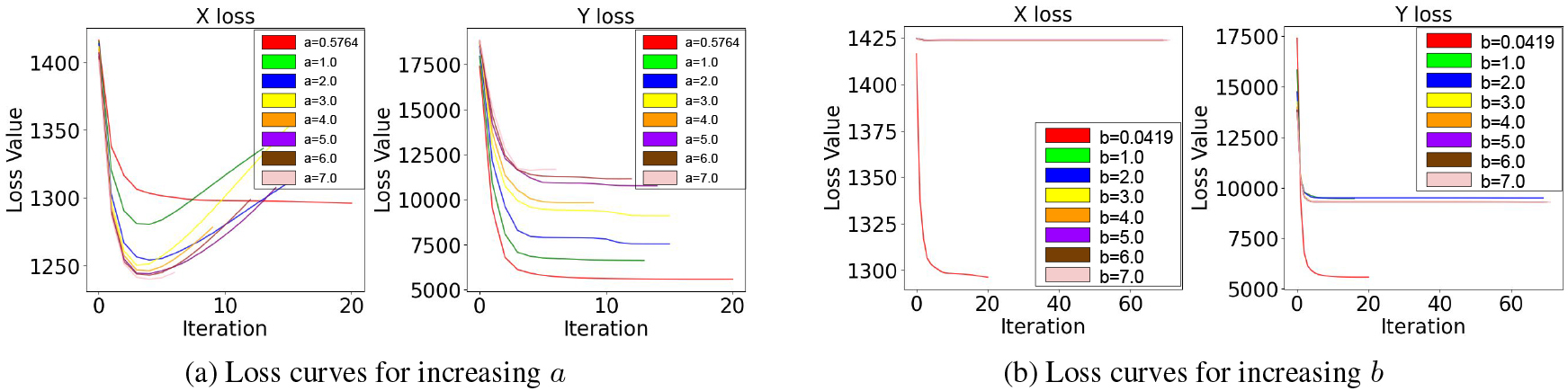
Model loss curves with increasing *a* (left) and *b* (right) weights.

#### 6.5.2 Impact of Rank on Performance

To evaluate the effect of rank on model performance, we conducted experiments using all methods with rank settings of {10, 20, 30, 40, 50, 60}. The corresponding results, measured by AUPR and AUC of both tensors *𝒳* and 𝒴, are presented in Figure 5. As shown in the two sub-figures on the bottom row, all models exhibit improved performance on the DDI prediction task (tensor 𝒴) as the rank increases, which is expected as higher rank in the latent space can capture more information. However, for the drug-drug combination prediction task (tensor *𝒳*), only SI-ADMM and TF-AI show improved performance as the rank increases. The performance of TDRC and CTF-DDI fluctuate when the value of rank changes. The performance of CP even declines with higher ranks (the two sub-figures on the top row in Figure 5). This indicates that these models face challenges in handling tensor data with varying structures and sparsity levels, two primary features that differentiate tensors *𝒳* and 𝒴. Notably, SI-ADMM demonstrates consistent performance improvement across both tensors as the rank increases.

**Figure 5.**
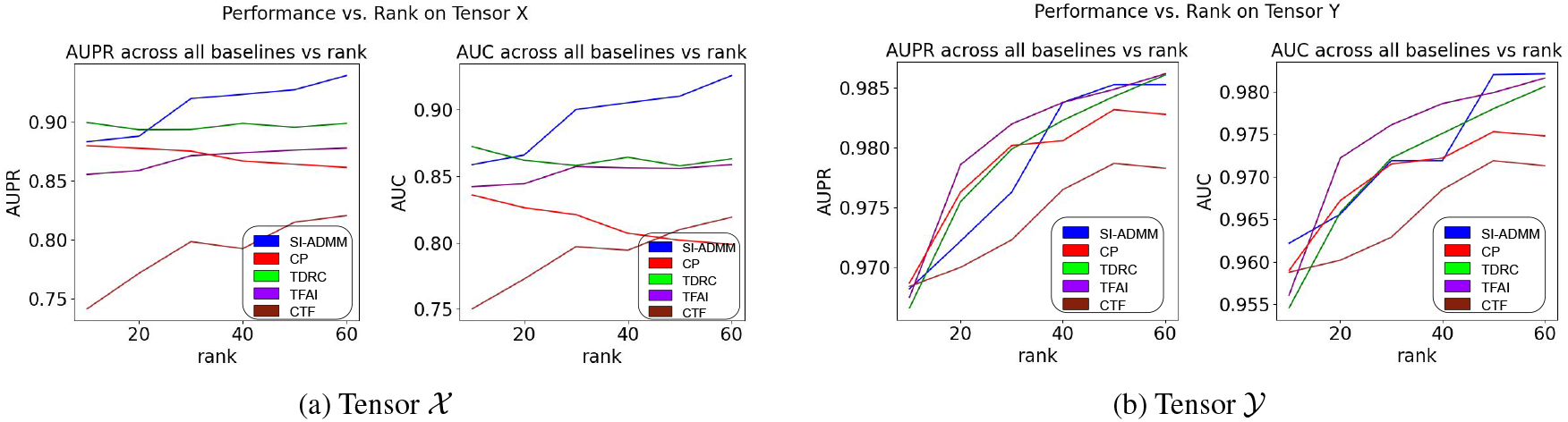
Model performance across ranks.

### 6.6 Runtime Analysis for SI-ADMM and Baselines

To evaluate the runtime of the SI-ADMM model, we conducted five independent runs for each method. The results are shown in figure 6. Since all models employ an early termination mechanism, we report the per-iteration runtime as a function of rank. For SI-ADMM in particular, the per-iteration runtime exhibits no significant increment as the rank grows. Accounting for early termination, the average total runtime of SI-ADMM still demonstrates no increase with respect to increasing rank. This relationship is consistent with the complexity analysis since the number of ranks (≤ 60) is much smaller than the number of drugs (1069). So the term that has the highest quantity becomes *N*^2^*R*(*M*_disease_ + *M*_DDI_), which makes the computation burden non-significant.

**Figure 6.**
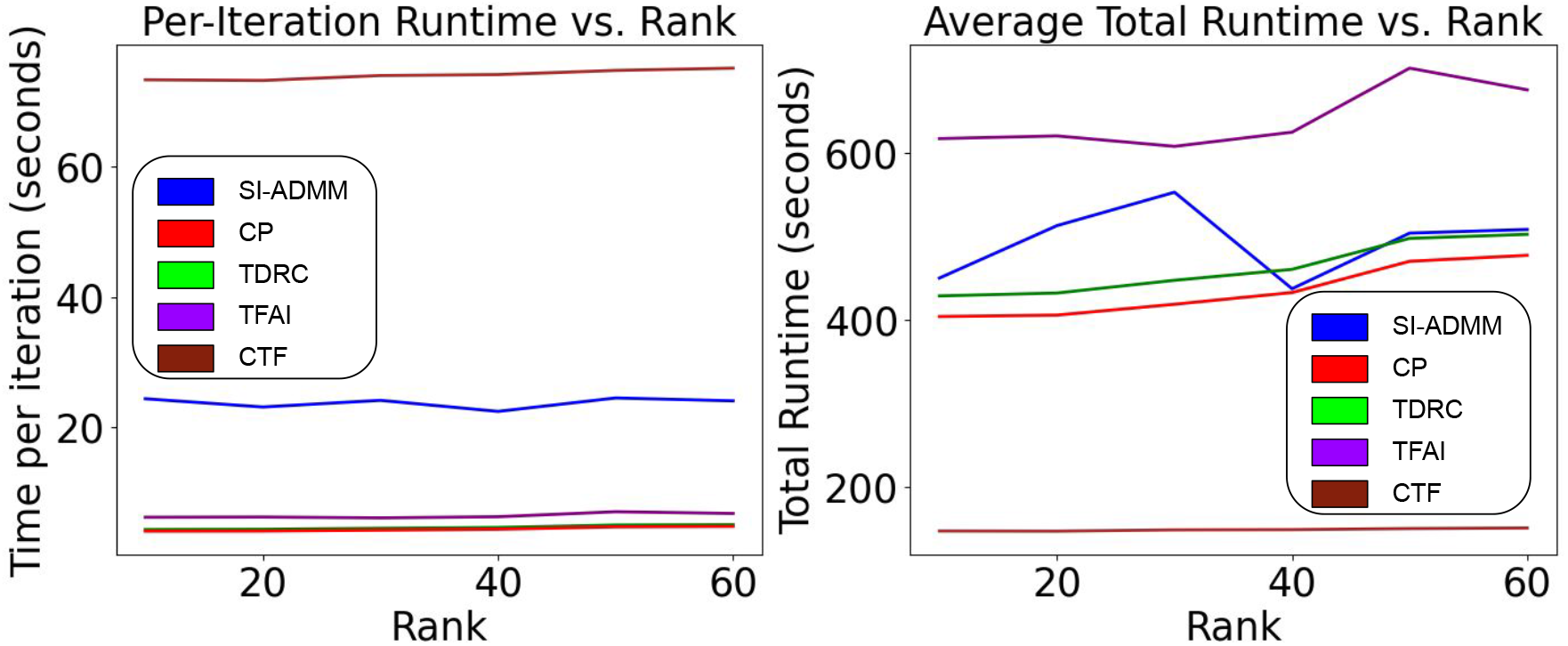
Runtime analysis of SI-ADMM and baselines. Left: per-iteration runtime (seconds) vs. rank. Right: average total runtime (five runs) vs. rank.

## 7 CONCLUSIONS

Drug-drug combinational therapies and drug-drug adverse effects are two closely related aspects of the same underlying phenomenon. In this paper, we proposed a joint tensor decomposition model, SI-ADMM, that simultaneously predicts disease-specific drug-drug combinations as well as DDIs, while mitigating sparsity through structured integration of auxiliary drug side information. We further developed an efficient optimization algorithm to solve the resulting joint tensor decomposition and reconstruction problem, and extensive experiments on real-world datasets demonstrated the effectiveness and robustness of the proposed SI-ADMM method. Looking ahead, our future work will focus on incorporating richer sources of side information and enhancing the model’s capacity for nonlinear transformations. In particular, we plan to integrate deep learning modules, such as graph neural networks (GNNs), to extract latent representations from molecular structures and heterogeneous biomedical networks [12, 27, 53]. In addition, to further improve expressive power, we intend to introduce neural network components after optimizing the latent factor tensors, allowing nonlinear transformations to be seamlessly incorporated while retaining the core optimization framework established in this work.

## 8 Acknowledgment

This work was supported by NSF CCF-2006780 and in part by NIH U01AG073323.

## Supplementary Appendix

This appendix provides the detailed derivations of the latent matrices *D, V, W, C*^(*i*)^, *U* ^(*i*)^

### Update of *D*

To calculate the partial derivative respect to *D* from equation (2), we can first separate out terms containing *D*.The terms in the Lagrangian ℒ that involve *D* are:

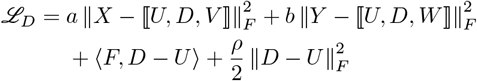

Using the approximation rule on the p.463 on the tensor review, the terms can be rewritten as:

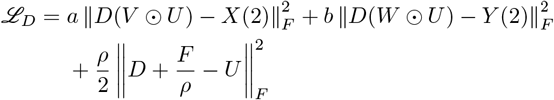

*X*(2), *Y* (2) are the second order matricizations of tensor X and Y. here set Φ = *V* ⊙*U* and Ψ = *W* ⊙ *U*, using equation 116 and 102 in Matrix Cookbook, the partial derivative of partial lagrangian function respects to *D* is:

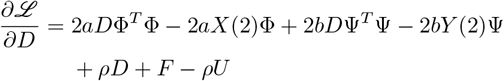

solve for *D* by setting the partial derivative to zero:

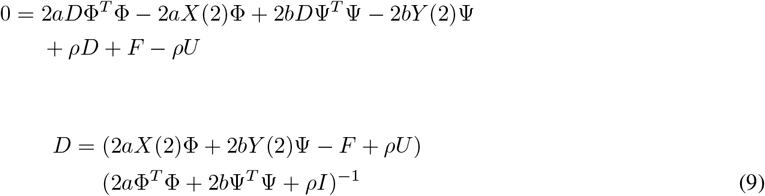

### Update of *V*

To calculate the partial derivative with respect to *V* from equation (2), we can first separate out terms containing *V*. The terms in the Lagrangian ℒ that involve *V* are:

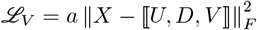

Using the approximation rule on the p.463 on the tensor review, the terms can be rewritten as:

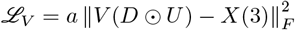

*X*(3) is the third order matricization of tensor X. Here set Ξ = *D* ⊙ *U*, using equations 116 and 102 in Matrix Cookbook, the partial derivative of partial lagrangian function respects to *V* is:

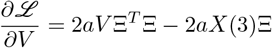

solve for *V* by setting the partial derivative to zero:

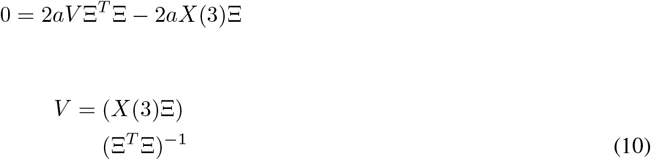

### Update of *W*

To calculate the partial derivative with respect to *W* from equation (2), we can first separate out terms containing *W*. The terms in the Lagrangian ℒ that involve *W* are:

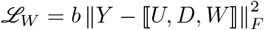

Using the approximation rule on the p.463 on the tensor review, the terms can be rewritten as:

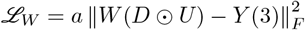

*Y* (3) is the third order matricization of tensor Y. Here set Ω = *D* ⊙ *U*, using equations 116 and 102 in Matrix Cookbook, the partial derivative of partial Lagrangianll function respects to *W* is:

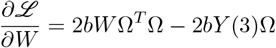

solve for *W* by setting the partial derivative to zero:

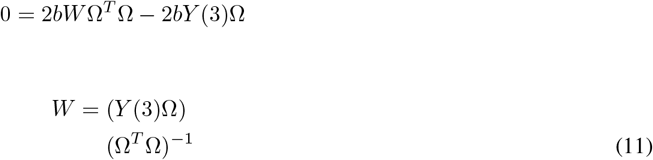

### Update of *U* ^(*i*)^ **and** *C*^(*i*)^

After the addition of the masking matrix on each side of the term in the second degree norm, the related terms for *U* ^(*i*)^ become:

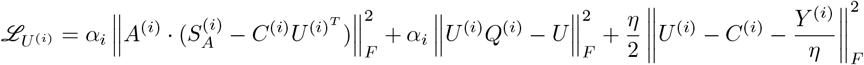

Here · is the element-wise multiplication, and *A*^(*i*)^ is a symmetric masking with values 1 for non-missing values and 0 for the missing value. The derivatives of the second and third terms are calculated in previous section, and the derivative of 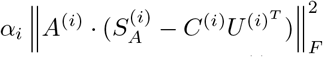 can be calculated as the following:

define matrix 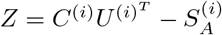, and Matrix 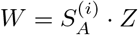, Then the term’s differential can be derived as:

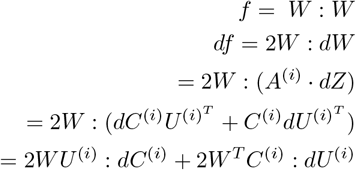

Here, : is the Frobenius norm product, and the partial derivative for *C*^(*i*)^ and *U* ^(*i*)^can be calculated as:

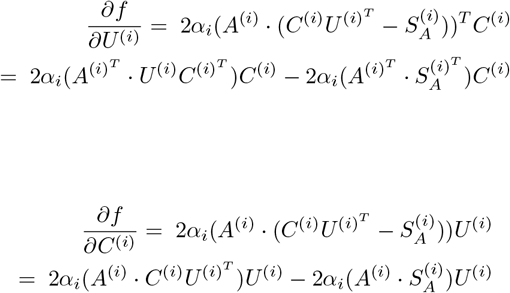

Putting the calculation of partial derivatives for all terms involves *C*^(*i*)^ and *U* ^(*i*)^ together, we got the over all partial derivative of the lagrangian function for the two matrices as:

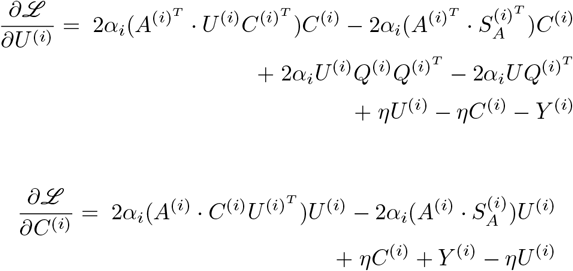

Since it is challenging to derive the analytic solve for *U* ^(*i*)^ and *U* ^(*i*)^ that separates them from the Hadamard product, we uses gradient descent to update *U* ^(*i*)^:

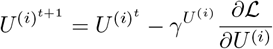

and use the similar gradient descent to update *C*^(*i*)^:

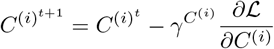

Here 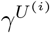 and 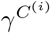 are the learning rates.

## Footnotes

1 https://icc.ise.bgu.ac.il/medical_ai/CDCDB/

2 https://tdcommons.ai/multi_pred_tasks/ddi

3 https://clinicaltrials.gov/

4 https://www.accessdata.fda.gov/scripts/cder/ob/

5 https://icd.who.int/icdapi

6 https://tdcommons.ai/multi_pred_tasks/ddi

## Notes

https://github.com/Xiaoge-Zhang/SI-ADMM

### Competing Interest Statement

The authors have declared no competing interest.

### Summary of Updates

Addition of Acknowledgment Section. Funding information is added to the manuscript. All other sections remain unchanged.

